# Motor sequence learning elicits mu peak-specific corticospinal plasticity

**DOI:** 10.1101/2024.07.31.606022

**Authors:** Tharan Suresh, Fumiaki Iwane, Minsu Zhang, Margaret McElmurry, Muskan Manesiya, Michael V Freedberg, Sara J Hussain

## Abstract

Motor cortical (M1) transcranial magnetic stimulation (TMS) interventions increase corticospinal output and improve motor learning when delivered during sensorimotor mu rhythm trough but not peak phases, suggesting that the mechanisms supporting motor learning may be most active during mu trough phases. Based on these findings, we predicted that motor sequence learning-related corticospinal plasticity would be most evident when measured during mu trough phases. Healthy adults were assigned to either a sequence or no-sequence group. Participants in the sequence group practiced the implicit serial reaction time task (SRTT), which contained an embedded, repeating 12-item sequence. Participants in the no-sequence group practiced a version of the SRTT that contained no sequence. We measured mu phase-independent and mu phase-dependent MEP amplitudes using EEG-informed single-pulse TMS before, immediately after, and 30 minutes after the SRTT in both groups. All participants performed a retention test one hour after SRTT acquisition. In both groups, mu phase-independent MEP amplitudes increased following SRTT acquisition, but the pattern of mu phase-dependent MEP amplitude changes after SRTT acquisition differed between groups. Relative to the no-sequence group, the sequence group showed greater peak-specific MEP amplitude increases 30 minutes after SRTT acquisition. Further, the magnitude of these peak-specific MEP amplitude increases was negatively associated with the magnitude of sequence-specific learning. Contrary to our original hypothesis, results revealed that motor sequence-specific learning elicits peak-specific corticospinal plasticity. Findings provide first direct evidence for the presence of a mu phase-dependent motor learning mechanism in the human brain.

**New and Noteworthy:** Recent work suggests that motor learning’s neural mechanisms may be most active during specific sensorimotor mu rhythm phases. If so, motor sequence learning-induced corticospinal plasticity should be more evident during some mu phases than others. Our results show that motor sequence-specific learning elicits corticospinal plasticity that is most prominent during mu peak phases. Further, this peak-specific plasticity correlates with learning. Findings establish the presence of a mu phase-dependent motor learning mechanism in the human brain.

## Introduction

The sensorimotor cortex is a hub within the human motor skill acquisition network. Human sensorimotor cortical activity dynamically oscillates in the mu (8-13 Hz) frequency range, with distinct phases of this rhythm reflecting brief windows of relative excitation and inhibition (Bergmann et al., 2019; Hussain et al., 2019; Suresh & Hussain, 2023; Zrenner et al., 2018). In non-human primates, motor cortical (M1) neuronal spiking rates are highest during mu rhythm trough phases and lowest during mu rhythm peak phases (Haegens et al., 2011). In humans, corticospinal output (Bergmann et al., 2019; Hussain et al., 2019; Suresh & Hussain, 2023; Wischnewski et al., 2022; Zrenner et al., 2018) and interhemispheric communication between homologous M1 subregions (Stefanou et al., 2018) are also increased during sensorimotor mu rhythm trough relative to peak phases.

In addition to single-neuron spiking rates, corticospinal output, and inter-hemispheric communication, sensorimotor mu rhythm phase covaries with the likelihood of inducing long-term potentiation-like (LTP-like) corticospinal plasticity in humans. For example, transcranial magnetic stimulation (TMS) interventions applied to M1 during mu rhythm trough phases preferentially induce LTP-like corticospinal plasticity, measured as increases in corticospinal output (Baur et al., 2020; Zrenner et al., 2018). In contrast, identical TMS interventions applied during mu rhythm peak phases induce weak long-term depression-like (LTD-like) corticospinal plasticity, measured as decreases in corticospinal output (Baur et al., 2020; Zrenner et al., 2018). These findings suggest that corticospinal plasticity induction is causally related to sensorimotor mu rhythm phase, with LTP-like and weak LTD-like corticospinal plasticity occurring preferentially during mu rhythm trough and peak phases, respectively. Given that LTP-like plasticity processes have been proposed to support human motor learning (Cantarero et al., 2013a; Cantarero et al., 2013b), and TMS interventions preferentially enhance motor learning when applied during mu rhythm trough phases (Hussain et al., 2021), the neurophysiological mechanisms underlying human motor learning may thus be most active during sensorimotor mu rhythm trough phases.

Outside of the motor system, brain oscillatory activity rhythmically controls various attentional (Busch & VanRullen, 2010; VanRullen et al., 2011), perceptual (Baumgarten et al., 2015; Busch et al., 2009; Dugué et al., 2011), and memory processes (Kerrén et al., 2018; Siegel et al., 2009). For example, the formation and retrieval of declarative memories has been proposed to occur during opposing hippocampal oscillatory phases (Hasselmo et al., 2002; Kerrén et al., 2018), object memory formation preferentially occurs during distinct prefrontal oscillatory phases (Siegel et al., 2009), and replay of whole-brain neuronal activity patterns associated with working memory are phase-locked to the theta rhythm (Fuentemilla et al., 2010). These findings suggest that oscillatory phase-dependent memory processing may be a widespread learning mechanism present across neural domains (Watrous et al., 2015). If so, phase-dependent neurophysiological signatures of motor learning should also be detectable within the human motor system.

Here, we reasoned that, if motor learning relies on a sensorimotor mu phase-dependent mechanism, then learning-related corticospinal plasticity should be more evident during some mu rhythm phases than others. Based on previous studies showing that corticospinal output (Hussain et al., 2019; Suresh & Hussain, 2023; Zrenner et al., 2018), interhemispheric communication (Stefanou et al., 2018), corticospinal plasticity (Baur et al., 2020; Zrenner et al., 2018), and the effects of TMS interventions on motor sequence learning (Hussain et al., 2021) are all enhanced during mu rhythm trough relative to peak phases, we predicted that motor sequence learning-related changes in corticospinal output would be more evident when measured during mu rhythm trough than peak phases.

To test this possibility, we measured sensorimotor mu phase-independent and phase-dependent corticospinal output using single-pulse, real-time EEG-informed TMS immediately before and after healthy adults practiced a serial reaction time task (SRTT) that included either an embedded, repeating 12-item motor sequence (sequence group) or a variation of the same task that included no sequence (no-sequence group). We predicted that mu phase-dependent corticospinal plasticity would be more prominent for the sequence group than the no-sequence group, with motor sequence-specific learning-related corticospinal plasticity being stronger when measured during mu trough than peak phases. As expected, our results show that participants in the sequence group exhibited greater sequence-specific learning than those in the no-sequence group. While both groups showed increased mu phase-independent corticospinal output following SRTT performance, each group showed a distinct pattern of mu phase-dependent corticospinal plasticity. Relative to the no-sequence group, the sequence group showed greater increases in *peak*-specific corticospinal output. Further, peak-specific increases in corticospinal output were negatively associated with sequence-specific learning. Surprisingly, these findings indicate that corticospinal plasticity induced by motor sequence-specific learning is more prominent during mu rhythm peak than trough phases. Findings establish the mechanistic role of mu rhythm phase in behaviorally-relevant corticospinal plasticity within the human motor system.

## Methods

### Participants

Thirty-six naïve right-handed healthy adult participants (age = 20.9 ± 0.52, 25 F and 11 M) participated in a single-session study that involved single-pulse transcranial magnetic stimulation (TMS), electroencephalography (EEG), electromyography (EMG), and behavioral testing. Right-handedness was confirmed using the Edinburgh Handedness Inventory (Oldfield, 1971). This study was approved by the Institutional Review Board at the University of Texas at Austin and all participants provided their written informed consent. Participants were randomly assigned to either the sequence group (N = 18, age = 20 ± 0.37 years, 13 F and 5 M) or the no-sequence group (N=18, age = 21.8 ± 0.94 years, 12 F and 6 M) prior to beginning experimental procedures. This sample size was chosen based on previous studies that identified significant mu phase-dependent variation in corticospinal output and corticospinal plasticity, as well as phase-dependent enhancement of motor sequence learning (N = 12-23, Zrenner et al., 2018; Bergmann et al., 2019; Hussain et al., 2019; Suresh & Hussain, 2023; Hussain et al., 2021).

### Experimental design and timeline

After EEG and EMG preparation, the scalp hotspot for the right first dorsal interosseous muscle (FDI) and the stimulation intensity needed to elicit motor-evoked potentials (MEPs) within this muscle were empirically determined for each participant. Participants then performed a choice reaction time task (cRT), after which parameters for real-time targeting of sensorimotor mu rhythm phases were individually optimized. Phase targeting accuracy was assessed during EEG recordings while participants rested quietly with their eyes open. Then, single-pulse TMS was delivered over the scalp hotspot during sensorimotor mu rhythm peak, trough, and random phases. Participants subsequently practiced one of two versions of the serial reaction time task (SRTT; see below for details). Immediately and 30 minutes after task completion, single-pulse TMS was again delivered during mu peak, trough and random phases. One hour after completing the SRTT, both groups performed a brief retention test that involved the same SRTT version performed during practice. After retention testing, all participants completed the Process Dissociation Procedure (PDP; Wilkinson & Shanks, 2004). Given that corticospinal plasticity following motor sequence learning under implicit and explicit task conditions may differ (Tunovic et al., 2014), the PDP was used to evaluate the development of explicit sequence awareness. Data from the cRT are not reported in this manuscript. See Figure 1 for experimental design and timeline.

**Figure 1.**
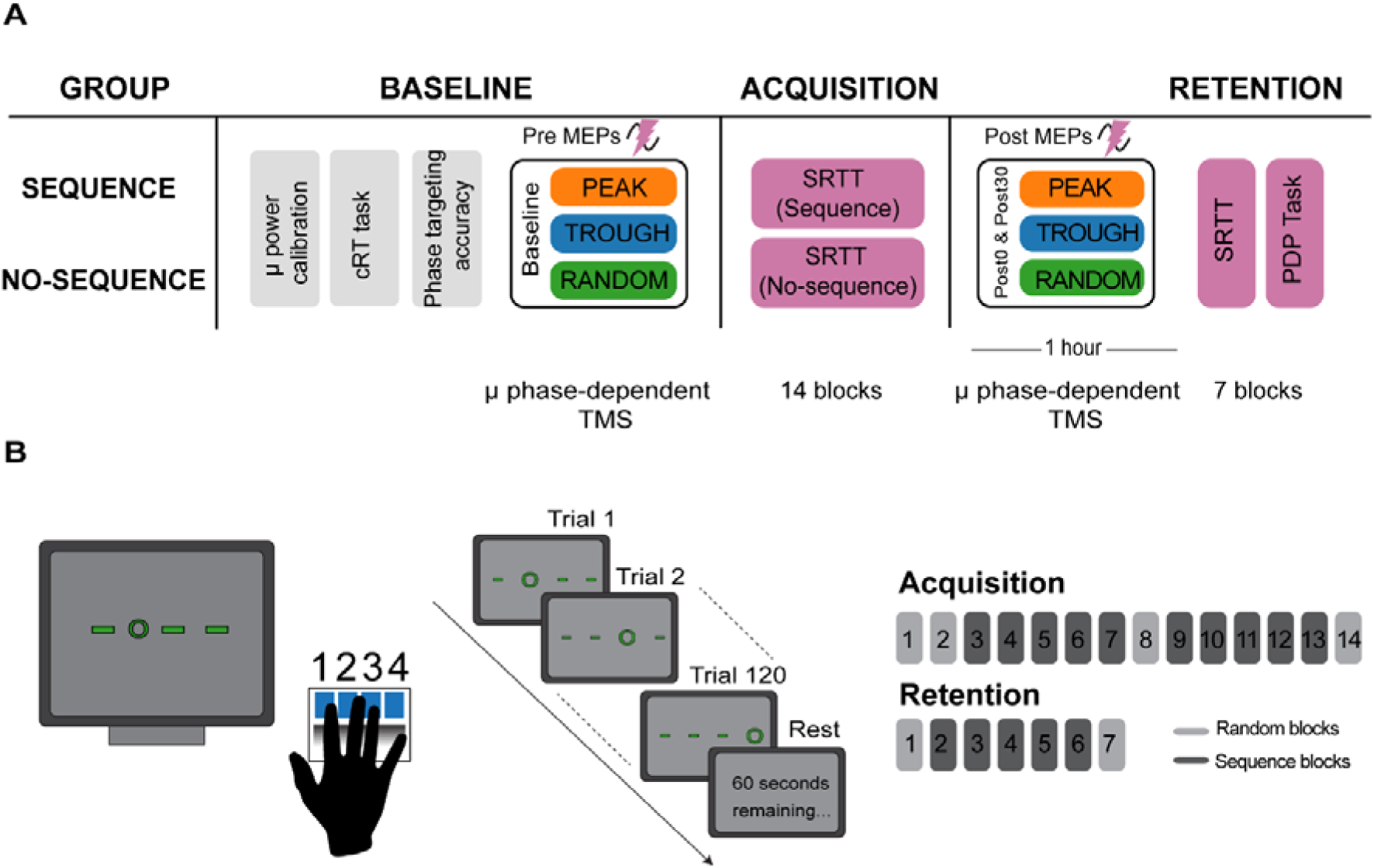
Experimental timeline and serial reaction time task. **A)** Experimental timeline. Real-time EEG analysis algorithm parameters were individually optimized during resting EEG recordings. After acquisition of phase-independent (random) and phase-dependent (peak and trough) MEPs at baseline, participants in both groups performed the SRTT. Post-SRTT phase-independent and phase-dependent MEPs were recorded immediately and 30 minutes after SRTT acquisition. SRTT retention was evaluated 1 hour after SRTT acquisition. **B)** Depiction of implicit SRTT. During acquisition, participants completed 14 blocks (120 trials per block). During retention, participants completed 7 blocks (120 trials per block). During sequence blocks, the sequence group practiced the SRTT with an embedded 12-item sequence, while during these blocks the no-sequence group practiced the task with no sequence (all random blocks).

### Data acquisition

#### EEG and EMG recording

64-channel EEG and bipolar EMG signals were recorded using TMS-compatible amplifiers (NeurOne Tesla with real-time digital out option, Bittium Biosignals, Finland). EEG impedances were maintained below 10 kΩ. EMG signals were recorded from the right FDI using disposable adhesive electrodes placed in a belly-tendon montage. EEG and EMG were recorded at a sampling rate of 5 kHz, with low-pass hardware filtering at 1250 Hz and a resolution of 0.001 µV.

#### Transcranial magnetic stimulation

We empirically determined the motor hotspot as the scalp site at which suprathreshold, single-pulse TMS elicited a reliable, focal twitch within the right FDI as well as a peak-to-peak motor evoked potential (MEP) amplitude of at least 50 μV. After hotspot identification, resting motor threshold (RMT) was determined using an automatic threshold-tracking algorithm (Borckardt et al., 2006). On average, RMT was 64.3 ± 3.8% of maximum stimulator output for the sequence group and 61.2 ± 3.1% for the no-sequence group. All TMS procedures were performed using a figure-of-eight coil held at ∼45° relative to the midsagittal line (DuoMAG XT-100, Deymed Diagnostic, biphasic pulse shape). Coil position accuracy was monitored online using frameless neuronavigation (BrainSight, Rogue Research) during hotspot identification, RMT determination, and mu phase-independent and mu phase-dependent single-pulse TMS.

#### Real-time EEG analysis

Real-time EEG-triggered TMS was performed using an established method (Zrenner et al., 2018; Shirinpour et al., 2020; Hussain et al., 2021; Suresh & Hussain, 2023). The EEG system was configured to stream data from 5 pre-selected EEG channels overlying the left sensorimotor cortex (C3, FC3, FC5, CP3 and CP5) using a local wired network connection (packet rate = 1 kHz, 5 packets per sample) between the EEG system and a dedicated PC for signal processing (Microsoft Windows 10, 10 cores, Intel i9 processor, 16 GB RAM) using Lab Streaming Layer software (Kothe, 2014) in real-time.

Custom MATLAB scripts were used to receive EEG data, process it, and send TTL triggers to the TMS device upon detection of the desired mu rhythm phase. During data streaming, EEG data was first downsampled to 1 kHz. To attenuate the effects of volume conduction, the Hjorth transformation was applied by subtracting the mean of the surrounding electrodes (FC3, FC5, CP3 and CP5) from the center electrode (C3; Hjorth 1975). The resulting signal was then buffered to create overlapping 500 ms windows for real-time analysis. At each time step, the current data window was zero-phase bandpass filtered using a Finite Impulse Response (FIR) filter (8-13 Hz, order = 128) and 64 ms data segments at the beginning and end of the 500 ms window were removed to eliminate edge effects. The remaining 372 ms data window was then used to fit an autoregressive model (Yule-Walker method, order: 30) that forward predicted the bandpass filtered signal 128 ms. After forward prediction, the Hilbert transform was used to generate an estimated phase angle timeseries signal. The phase estimation algorithm was also configured to trigger TMS only when each participant’s sensorimotor mu power exceeded a predefined threshold. To calculate this threshold, resting state EEG data was divided into non-overlapping 500 ms segments, bandpass filtered using a Finite Impulse Response (FIR) filter (8-13 Hz, order = 128), Hjorth transformed, downsampled to 1 kHz, and then spectrally decomposed (Welch’s method, 4-100 Hz with 0.25 Hz resolution excluding frequencies between 57.75 – 62 Hz to reduce the influence of line noise). For each participant, the median of the resulting frequency-domain signal between 8 and 13 Hz was defined as the power threshold. When the estimated instantaneous phase angle at the time point of interest matched the pre-determined mu phase condition and mu power exceeded each participant’s threshold, a single TTL pulse was sent to the TMS system and also marked the time of TMS delivery in the EEG recording.

#### Phase targeting accuracy and real-time EEG-informed single-pulse TMS

Prior to delivering real-time EEG-informed TMS, we first evaluated the accuracy of phase targeting in the absence of TMS delivery in each participant. 25 TTL pulses were sent to the EEG system at mu peak, trough and random phases, but no TMS was delivered (minimum inter-stimulus interval = 5 s + random jitter). During mu phase-independent single-pulse TMS, blocks of 25 TMS pulses were delivered to the scalp hotspot for the right FDI during random mu phases (120% RMT, inter-stimulus interval = 5 s + random jitter). During mu phase-dependent single-pulse TMS, blocks of 25 TMS pulses were delivered during mu peak and trough phases (120% RMT, minimum inter-stimulus interval = 5 s + random jitter). Real-time EEG-informed single-pulse TMS was delivered at baseline, immediately after SRTT acquisition, and 30 minutes after SRTT acquisition. The order of phases targeted per block (i.e., random, peak, and trough) was counterbalanced across participants in both groups.

#### Serial Reaction Time Task

Participants in both the sequence and no-sequence groups were seated in front of a computer monitor with their right arm resting on a tabletop and their right hand positioned atop a serial response box (Chronos Response Device, Psychology Software Tools, see Figure 1). Participants performed one of two versions (sequence or no-sequence) of the SRTT (Nissen & Bullemer, 1987; Perez et al., 2007) programmed using E-prime 3 (Psychology Software Tools). The no-sequence group was included in this study to experimentally account for non-specific effects of motor task engagement (including repeated keypresses) on MEP amplitude changes. During the SRTT, the monitor displayed four equally-spaced green dashes against a black background. On each trial, one of these dashes was replaced by a circle. Prior to beginning the task, participants were instructed to respond to the presentation of the circle as fast and accurately as possible by pressing one of the four keys with the corresponding finger of their right hand, as shown in Figure 1. When the participant pressed the correct key, the circle disappeared, and the trial was complete. If an incorrect key was pressed, the circle remained on the monitor and the trial continued until the correct key was pressed. 500 ms elapsed between trials. No additional feedback on keypress accuracy or keypress latency was provided.

Participants became familiarized with the task during a practice block that included 10 trials. During SRTT acquisition, participants in the sequence group performed 14 blocks of keypresses, with one minute of rest between blocks. Each block included 10 repetitions of a 12-item sequence, amounting to a total of 120 keypresses per block. Blocks 3-7 and 9-13 included a pre-determined, embedded 12-item sequence (2-3-1-4-3-2-4-1-3-4-2-1). Blocks 1, 2, 8 and 14 were random blocks that contained no embedded sequence. During SRTT retention, participants in the sequence group performed 7 blocks of 120 keypresses each, again resting for one minute between blocks. Blocks 1 and 7 were random blocks while blocks 2-6 were sequence blocks.

Random blocks contained no consecutive item repeats and each item had approximately the same frequency of appearance. Participants in the no-sequence group performed randomized keypresses for all blocks. Participants were not informed about the presence or absence of the sequence or its length.

#### Process Dissociation Procedure

After SRTT retention, participants completed the PDP. The PDP assesses conscious control over sequence knowledge that may develop during implicit SRTT performance (Destrebecqz et al., 2005; Wilkinson et al., 2009, 2015; Wilkinson & Jahanshahi, 2007; Wilkinson & Shanks, 2004) and is based on the rationale that conscious knowledge is both reportable upon request and under intentional control. The PDP consists of an inclusion and exclusion test, each of which involve 12 unique three-item fragments of the 12-item sequence that were embedded within the sequence version of the SRTT (e.g., 2-3-1, 3-1-4, 1-4-3, etc.).

The inclusion test evaluates reportable, consciously accessible knowledge. During the inclusion test, participants were presented with two elements of each of 12 three-item fragments and were asked to produce the next element in the fragment by pressing the corresponding button. The exclusion test evaluates intentional control. During the exclusion test, participants were again presented with two elements of each of 12 three-item fragments and were asked to *avoid* producing the next element in the fragment by pressing any button *other than* the one that corresponded to the next element in the fragment. At the beginning of the inclusion and exclusion tests, participants were informed that repeated element sequences (i.e., a 2 immediately after a 2) were not allowed. For each trial of the inclusion test, there was a 1/3 probability of pressing the correct key by chance. For each trial of the exclusion test, there was a 2/3 probability of pressing the correct key by chance. The order of inclusion and exclusion tests was counterbalanced across participants in both groups.

### Data analysis

#### Serial Reaction Time Task

Reaction times (RTs) were averaged within each block. We defined acquisition and retention skill as the difference between the mean RT of the last random block and last sequence block during SRTT acquisition and the difference between the mean RT of the first random block and first sequence block during SRTT retention, respectively. We also defined offline changes in skill as the difference between acquisition skill and retention skill. Here, positive values reflect offline losses while negative values reflect offline gains.

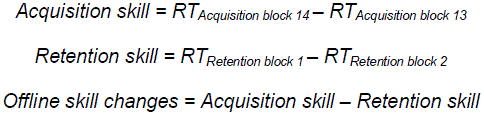

#### Process Dissociation Procedure

Keypresses during inclusion trials were considered accurate when the participant correctly completed the 3-element fragment by pressing the correct key in the sequence. Keypresses during exclusion trials were considered accurate when the participant correctly avoided the next element in the fragment by pressing one of the two other key choices. The total number of accurate trials in the inclusion and exclusion conditions were compared to the corresponding chance levels (i.e., 4/12 for the inclusion test and 8/12 for the exclusion test) to test for explicit awareness of the embedded sequence.

#### EEG analysis

EEG signals were processed using a combination of Fieldtrip (Oostenveld et al., 2010) and custom-written MATLAB scripts. To evaluate phase targeting accuracy, we segmented EEG data into 6 s segments (± 3 s relative to the targeted phase), and then demeaned and detrended these segments. To ensure that all offline data analysis procedures mimicked those applied during real-time phase targeting, we applied the Hjorth transformation to the EEG data by subtracting the mean of four surrounding sensorimotor electrodes (FC3, FC5, CP3 and CP5) from the center sensorimotor electrode (C3). Resulting data segments were zero-phase bandpass filtered using an FIR filter (8-13 Hz). The instantaneous phase of the filtered signal at the time of phase targeting was identified using the Hilbert transformation. We then combined phase angles across all participants per group.

Given that mu phase-power interactions shape MEP amplitude (Hussain et al., 2019; Ozdemir et al., 2022; Suresh & Hussain, 2023), we also calculated pre-stimulus sensorimotor mu rhythm power. Continuous EEG data obtained during TMS delivery were segmented into 500 ms segments (0.51 to 0.01 seconds prior to each TMS pulse), demeaned, linearly detrended and Hjorth transformed (center = C3, surround = FC1, FC5, CP1, CP5; Hjorth, 1975; Zrenner et al., 2018; Hussain et al., 2021; Suresh & Hussain, 2023). The resulting Hjorth-transformed signals were then spectrally decomposed (Welch’s method, 4-50 Hz with 0.25 Hz resolution, Suresh & Hussain, 2023). Each trial’s power values were averaged in the mu (8-13 Hz) range to generate a mixed mu power estimate (i.e., the combination of aperiodic and periodic components of mu power; Donoghue et al., 2020). Mixed mu power was calculated instead of periodic mu power as our previous work demonstrated that statistical models including mu rhythm phase and mixed mu power explain MEP amplitude variation better than those including mu phase and periodic mu power (Suresh & Hussain, 2023).

#### MEP analysis

Pre-stimulus EMG activity was filtered with a discrete Fourier transform filter to reduce line noise. All trials were visually inspected, and those that precluded reliable calculation of peak-to-peak MEP amplitudes were rejected. We also calculated background EMG as the root-mean-square of pre-stimulus EMG activity between 100 and 25 ms before each TMS pulse; trials with values greater than 10 µV were rejected. Peak-to-peak MEP amplitudes were calculated as the maximum voltage deflection between 20 and 40 ms after each TMS pulse. Overall, 5.5% of all trials were rejected. MEP amplitudes were log-transformed to reduce positive skew and improve statistical model fits (Hussain et al., 2019; Suresh & Hussain, 2023).

### Statistical analysis

#### Phase targeting accuracy

Phase angle distributions at the time of phase targeting were evaluated for deviations from a circular uniform distribution in the 90° (peak phases) and 270° (trough phases) directions using separate V-tests for each group. For random phases, general deviations from circular uniformity were tested using Rayleigh’s test. For each group, mu peak and trough phase angle distributions were compared using the Watson-Williams test. Phase angle distributions for each condition (peak, trough, and random) were also compared across groups using the Watson-Williams test.

#### SRTT acquisition and retention

To evaluate changes in mean RT during SRTT acquisition and retention, we fit separate linear mixed-effects models (LMMs) for data acquired during SRTT acquisition and retention periods. These included mean RT as the response variable, BLOCK (categorical variable: acquisition blocks 1-14, retention blocks 1-7), GROUP (categorical variable: sequence, no-sequence), and BLOCK x GROUP interactions as fixed effects. PARTICIPANT was included as random intercept. To evaluate changes in skill from SRTT acquisition to retention, we fit an LMM that included the change in mean RT as the response variable, TASK (categorical variable: acquisition, retention, offline change), GROUP (categorical variable: sequence, no-sequence), and TASK x GROUP interactions as fixed effects. Post hoc pairwise comparisons were used to compare SRTT acquisition and retention between groups. To evaluate offline skill, we compared offline changes to zero using separate single-sample t-tests for the sequence and no-sequence groups.

#### Process dissociation procedure (PDP)

To assess explicit sequence awareness, we compared fragment completion accuracy to chance using separate single-sample right-tailed t-tests for the inclusion and exclusion tests. One participant in the no-sequence group did not complete the PDP.

#### MEP analysis

We first examined the influence of SRTT acquisition on phase-independent MEP amplitudes using an LMM with GROUP (categorical variable: sequence, no-sequence), TIME (categorical variable: baseline, immediately after SRTT acquisition, 30 minutes after SRTT acquisition), and the GROUP x TIME interaction as fixed factors and natural log-transformed POWER (continuous variable) as a covariate, and natural log-transformed MEP amplitudes elicited during random mu phases as the response variable. The random intercept of PARTICIPANT was also included (Hussain et al., 2019; Hussain et al., 2021; Suresh & Hussain, 2023). Post hoc pairwise comparisons were used to compare phase-independent MEP amplitudes across time points.

Next, we evaluated the influence of SRTT task performance on phase-dependent MEP amplitudes by fitting an LMM that included fixed effects of GROUP (categorical variable: sequence, no-sequence), PHASE (categorical variable: peak, trough), TIME (categorical variable: baseline, immediately after SRTT acquisition, 30 minutes after SRTT acquisition), as well as their two– and three-way interactions as fixed factors, the PHASE x natural log-transformed POWER interaction as a covariate, and natural log-transformed MEP amplitudes as the response variable. PARTICIPANT was included as the random intercept. This LMM revealed a significant three-way interaction (see *Results*). To further characterize this three-way interaction, we fit separate LMMs for each group. These post hoc LMMs included PHASE (categorical variable: peak, trough), TIME (categorical variable: baseline, 30 minutes after SRTT acquisition), and the PHASE x TIME interaction as fixed factors, as well as the PHASE x log-transformed POWER interaction as a covariate. Natural log-transformed mu MEP amplitudes were the response variable, and PARTICIPANT was included as the random intercept. Post hoc pairwise comparisons were used to compare MEP amplitudes across time points and phases. Given the mu PHASE x POWER interaction identified in our previous work and that of others (Bigoni et al., 2024; Hussain et al., 2019; Ozdemir et al., 2022; Suresh & Hussain, 2023) and our primary interest in evaluating mu phase-dependent MEP amplitude changes, we included the PHASE x natural log-transformed POWER interaction as a covariate in all our LMMs to control for this known interaction effect (Hussain et al., 2019; Suresh & Hussain, 2023).

Finally, we examined relationships between phase-dependent MEP amplitude changes and SRTT performance changes. We first used the LMM that evaluated phase-dependent changes in MEP amplitudes across groups to obtain trial wise peak– and trough-specific model-fitted MEP amplitudes at baseline and 30 minutes after SRTT acquisition. These time points were chosen because our phase-dependent MEP analysis revealed a significant three-way GROUP x PHASE x TIME interaction across these two time points (see *Results*), Then, we calculated the magnitude of phase-dependent MEP amplitude changes for each participant by dividing the fitted mean phase-dependent MEP amplitudes 30 minutes after SRTT acquisition by the fitted mean phase-dependent MEP amplitudes at baseline.

Separate LMMs were then used to evaluate relationships between (1) mu peak-specific MEP changes and SRTT performance and (2) mu trough-specific MEP changes and SRTT performance. Each LMM included GROUP (categorical variable: sequence, no-sequence), TASK (categorical variable: acquisition, retention), MEP ratio (continuous variable), and their two– and three-way interactions as fixed factors. Each LMM included also SKILL (i.e., change in mean RT during SRTT acquisition and retention, see *Data analysis*) as the response variable and PARTICIPANT as the random intercept. Post hoc pairwise comparisons were used to statistically evaluate relationships between peak-specific MEP amplitude changes and SRTT performance for each group.

All statistical analyses were performed in R (version 4.3.1). Model fits were visually inspected using histograms of residuals and quantile-quantile plots. The significance of fixed effects was determined using likelihood ratio tests. All reported β-coefficients represent the difference in estimates between groups and phases. Alpha was equal to 0.05 for all comparisons and p-values were corrected using the Tukey’s Honestly Significant Difference test for multiple comparisons where appropriate. Data are expressed as mean ± standard error of the mean.

## Results

### Phase targeting accuracy

Real-time EEG analysis accurately identified and targeted mu peak (Figure 2, target phase = 90°, observed mean phase for sequence group = 85.7° [95% confidence interval {CI} = 80.27°– 91.12°], V = 237.9, p < 0.001; observed mean phase for no-sequence group = 86.93° [95% CI = 80.77°– 93.10°], V = 217.52, p < 0.001) and mu trough phases (target phase = 270°, observed mean phase for sequence group = 271.72° [95% CI = 266.17° – 277.25°], V = 56.01, p < 0.001; observed mean phase for no-sequence group = 269.66° [95% CI = 263.99° – 275.31°], V = 45.71, p = 0.001). As expected, targeted phases differed significantly between peak and trough conditions in both the sequence group (F = 1687.0, p < 0.001) and the no-sequence group (F = 1495.8, p < 0.001). For random phases, we observed a deviation from circular uniformity in the 20° direction in the sequence group (R = 5.56, p = 0.003, observed mean phase = 19.05° [95% CI = 343.15° – 54.95°]) and no significant deviation from uniformity in the no-sequence group (R = 2.47, p = 0.08, observed mean phase = 12.93° [95% CI = 311.2° – 74.66°]). However, we observed no significant differences in phase targeting accuracy between the sequence and no-sequence groups for any phase (peak: F = 0.11, p = 0.741; trough: F = 0.341, p = 0.559; random: F = 0.188, p = 0.665).

**Figure 2.**
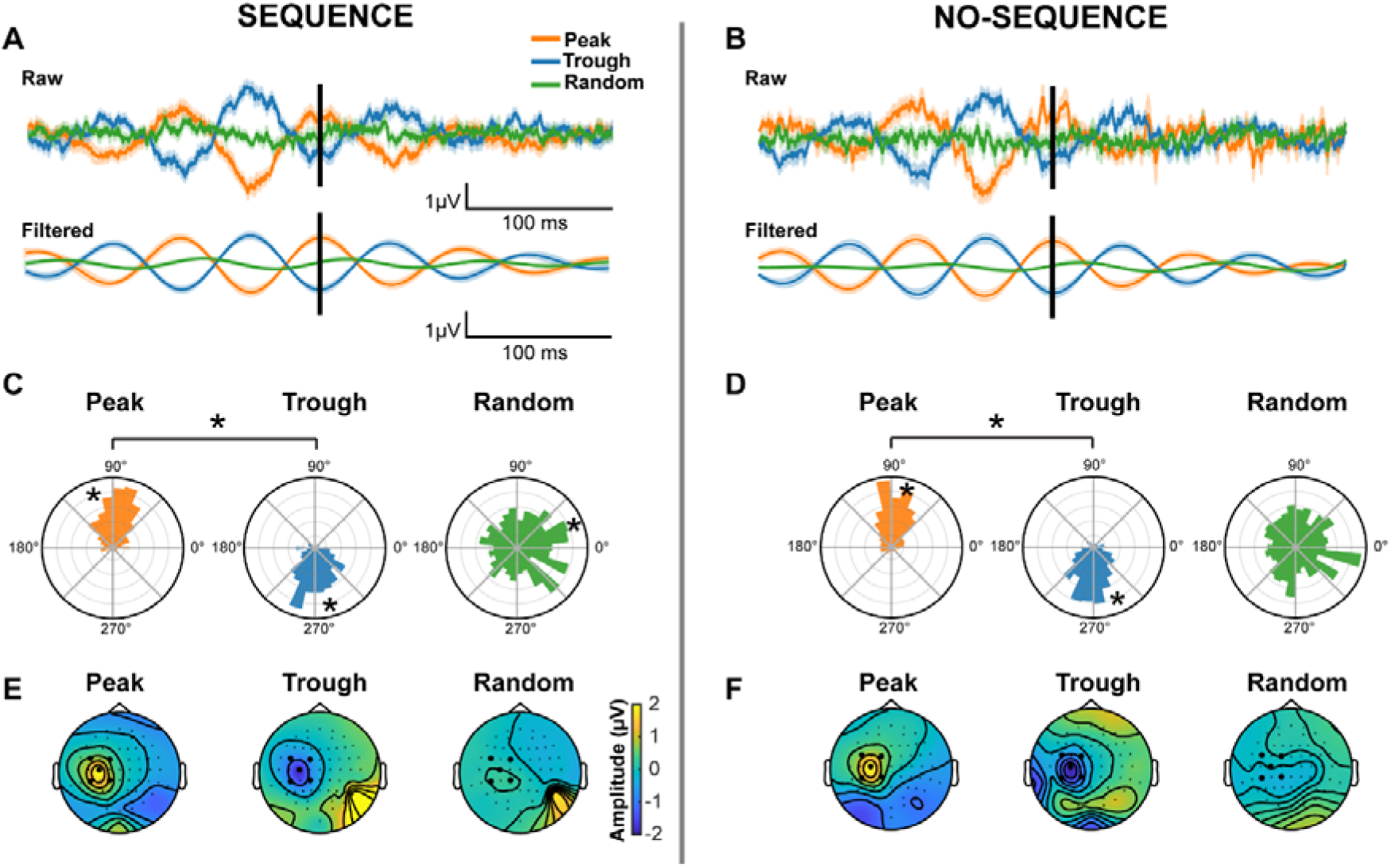
Phase targeting accuracy. **A, B)** Group-averaged raw and filtered (8-13 Hz) EEG signals recorded over left sensorimotor cortex during evaluation of mu phase targeting accuracy in the sequence (A) and no-sequence (B) groups. Black vertical lines reflect the time point at which the real-time EEG analysis algorithm identified mu peak, trough, and random phases. Shading represents mean ± 1 SEM. **C, D)** Circular histograms indicate the phase angle of the sensorimotor mu (8-13 Hz) rhythm at the time of phase targeting for peak, trough, and random phases in the sequence (C) and no-sequence (D) groups. **E, F)** Scalp distribution of C3-Hjorth transformed and band-pass filtered EEG activity during mu peak, trough and random phases at the time of phase targeting for each phase condition in the sequence (E) and no-sequence (F) groups. Topographical plots reflect average scalp-recorded EEG signals (filtered between 8-13 Hz) within a 5 ms window centered on the time of phase targeting. Bolded channels indicate those used to calculate C3-Hjorth transformed EEG data. Black asterisks reflect statistical significance.

### SRTT

During SRTT acquisition, participants in both the sequence and no-sequence groups decreased their mean RTs across blocks (Figure 3, main effect of BLOCK: F = 12.96 and p < 0.001). RTs did not differ between groups for random blocks, but participants in the sequence group tended to show faster mean RTs during the sequence blocks than the no-sequence group during the corresponding blocks (BLOCK x GROUP interaction: F = 9.37 and p < 0.001; no main effect of GROUP: F = 0.63 and p = 0.42). However, post hoc pairwise comparisons did not identify any significant differences in mean RTs between groups during these blocks (i.e., blocks 3-7 or 9-13, see Figure 3). For both groups, mean RTs were not statistically distinguishable between the first and last random blocks (blocks 1 vs. 14, t < 0.165, p = 0.99). For the sequence group, mean RTs were faster during the last sequence block than the last random block (block 13 vs. 14, t ratio = –8.51, p < 0.001), but this was not the case for the no-sequence group (t ratio = 0.16, p = 0.99). Finally, acquisition skill was significantly greater in the sequence than no-sequence group (Figure 3, t ratio = 5.94, p < 0.001). Thus, as expected, participants in the sequence group experienced sequence-specific learning.

**Figure 3.**
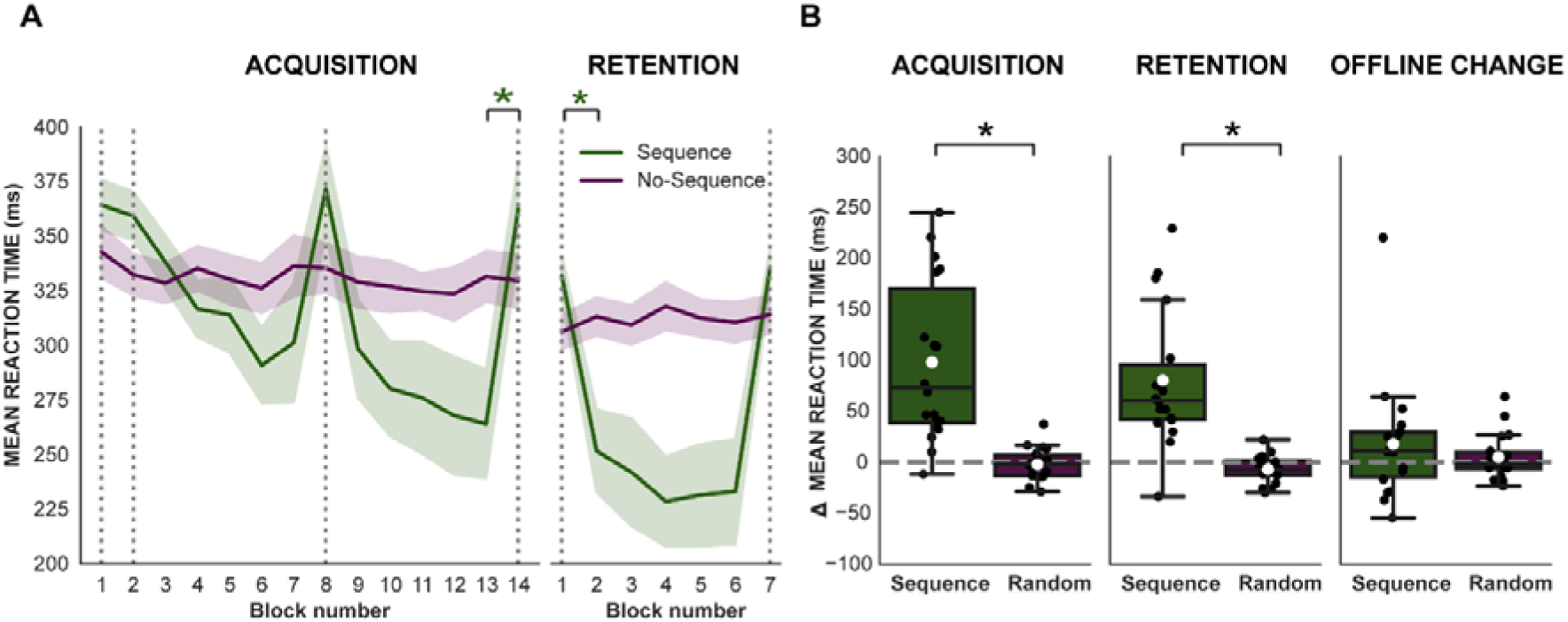
Serial reaction time task performance. **A)** Participants in the sequence group showed sequence-specific decreases in mean RTs during SRTT acquisition and retention. Vertical dotted lines indicate random blocks during the SRTT (practice block not shown). Green asterisks reflect significant post hoc pairwise comparisons between blocks 13 and 14 during SRTT acquisition and blocks 1 and 2 during retention for the sequence group. Shading represents the mean ± 1 SEM. **B)** Sequence-specific skill acquisition and retention were greater in the sequence group than in the no-sequence group. Black asterisks reflect significant post hoc comparisons for acquisition, retention and offline skill changes between groups. Note that for offline changes, positive values reflect offline losses while negative values reflect offline gains. For boxplots, black lines represent median, white dots represent mean, black dots represent individual participants, and error bars represent upper and lower inter-quartile ranges.

During SRTT retention, participants in both groups decreased their mean RTs across blocks (Figure 3, main effect of BLOCK: F = 18.91 and p < 0.001) and mean RTs were overall faster for the sequence group than the no-sequence group (main effect of GROUP: F = 6.04, p = 0.01). Mean RTs were not statistically distinguishable between groups during random blocks but were significantly faster for the sequence group than the no-sequence group during sequence blocks (GROUP x BLOCK interaction: F = 21.14, p < 0.001). Post hoc pairwise comparisons revealed that mean RTs were faster for the sequence group than for the no-sequence group during blocks 4-6 (t > 3.59, p values < 0.03). In addition, retention skill was significantly greater in the sequence group than the no-sequence group (Figure 3, t ratio = 5.17, p < 0.001). Thus, participants in the sequence group retained sequence-specific skill obtained during SRTT acquisition. Finally, participants in the sequence group showed significant offline losses (t = 5.23 p < 0.0001, mean = 98.17 ms [95% CI = 58.61 ms – 137.72 ms] while those in the no-sequence group did not (t = –0.51, p = 0.616, mean = –1.95 ms [95% CI = –10.01 ms – 6.11ms]). However, direct comparison of the magnitude of offline changes revealed no significant difference between groups (Figure 3, t = 0.76, p = 0.97).

### Explicit sequence awareness

In the sequence group, mean fragment completion accuracies exceeded chance for the inclusion test (6 ± 0.66 [SEM], t = 3.02, p = 0.007, chance = 4) but not the exclusion test (7.72 ± 0.54, t = –0.51, p = 0.61, chance = 8). In the no-sequence group, fragment completion accuracies did not exceed chance for either the inclusion or exclusion tests (Figure 4, inclusion = 3.88 ± 0.45, exclusion = 7.7 ± 0.54, t < –0.52 for both, p > 0.59 for both). Given that explicit awareness requires both conscious knowledge of the sequence and the ability to intentionally control sequence production, there was no strong evidence of explicit sequence awareness in either group.

**Figure 4.**
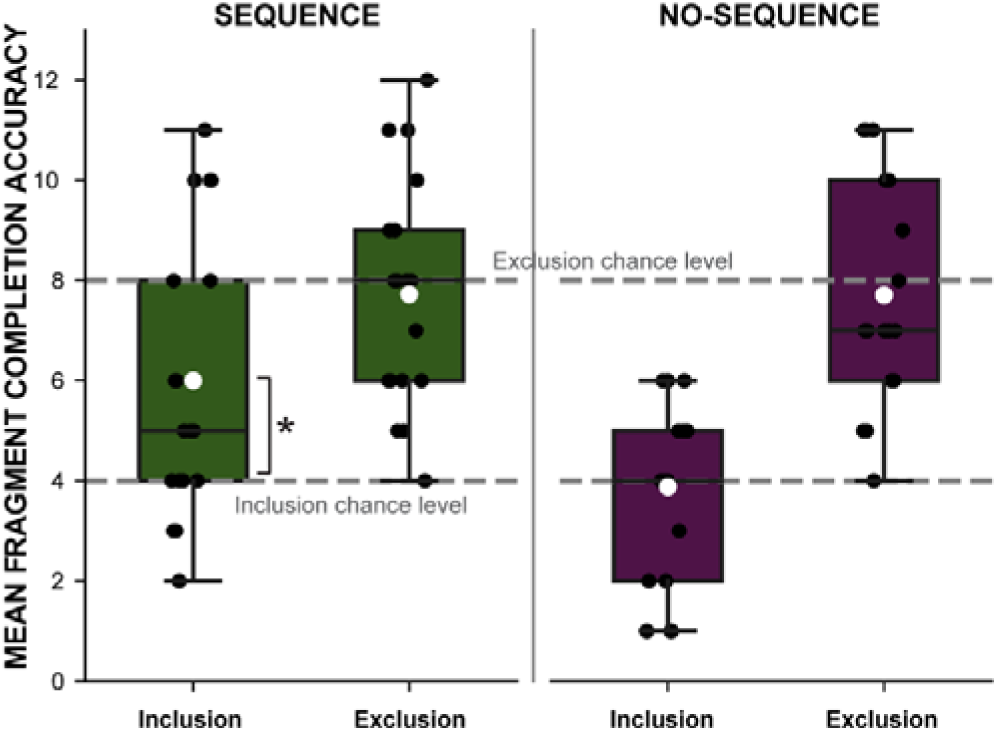
Process Dissociation Procedure. Mean number of fragments completed or avoided during the inclusion and exclusion portions of the PDP, respectively. The dashed lines reflect chance performance (4/12 for inclusion and 8/12 for exclusion). Black asterisks reflect significant difference between mean fragment completion accuracy and inclusion chance level in the sequence group. For boxplots, black lines represent group median, white dots represent group mean, black dots represent individual participants, and error bars represent upper and lower inter-quartile ranges.

### Phase-independent changes in MEP amplitudes

We first characterized the effects of SRTT acquisition on phase-independent MEP amplitudes elicited during random mu phases. In both groups, MEP amplitudes increased following SRTT acquisition (significant main effect of TIME; F = 30.01, p < 0.001, no main effect of GROUP; F = 0.68, p = 0.41, no GROUP x TIME interaction, F = 0.88, p = 0.41). Post hoc pairwise comparisons revealed that random, phase-independent MEP amplitudes increased from baseline to immediately after SRTT acquisition (β = –0.08, t ratio = –2.38, p = 0.04), from baseline to 30 minutes after SRTT acquisition (β = –0.25, t ratio = –7.58, p < 0.0001), and from immediately to 30 minutes after SRTT acquisition (β = –0.17, t ratio = – 5.21, p < 0.001). Thus, phase-independent MEP amplitudes increased similarly following SRTT acquisition in both groups.

### Phase-dependent changes in MEP amplitudes

We next evaluated the effects of SRTT acquisition on phase-dependent MEP amplitudes elicited during mu peak and trough phases. We found a significant three-way GROUP x TIME x PHASE interaction (Figure 5, likelihood ratio test: F = 4.95, p = 0.007). However, this three-way interaction was only present when comparing changes in MEP amplitudes from baseline to 30 minutes after SRTT acquisition (no GROUP x TIME x PHASE interaction for baseline versus immediately after SRTT: t = –0.9, p = 0.36, significant GROUP x TIME x PHASE interaction for baseline versus 30 minutes after SRTT: t = 2.19, p = 0.02), indicating that the magnitude of MEP amplitude changes from baseline to 30 minutes after SRTT acquisition more strongly differed across mu phases in one group than the other. Given the significant three-way GROUP x PHASE x TIME interaction present from baseline to 30 minutes after SRTT acquisition, we evaluated mu phase-dependent changes in MEP amplitudes across these two time points separately for each group. For the sequence group, this post hoc analysis revealed a significant effect of PHASE (F = 5.32, p = 0.02) and TIME (F = 68.59, p < 0.0001), but no PHASE x TIME interaction (F = 1.29, p = 0.25). Additional post hoc pairwise comparisons further revealed that MEP amplitudes were significantly larger during mu trough than peak phases (β = –0.08, t ratio = –2.35, p = 0.01), while MEP amplitudes elicited during both phases were significantly larger 30 minutes after SRTT acquisition than baseline (β = –0.29, t ratio = –8.28, p < 0.0001). For the no-sequence group, post hoc analysis revealed a significant effect of PHASE (F = 24.9, p < 0.0001), TIME (F = 104.89, p < 0.0001) and a significant PHASE x TIME interaction (F = 4.06, p = 0.04).

**Figure 5.**
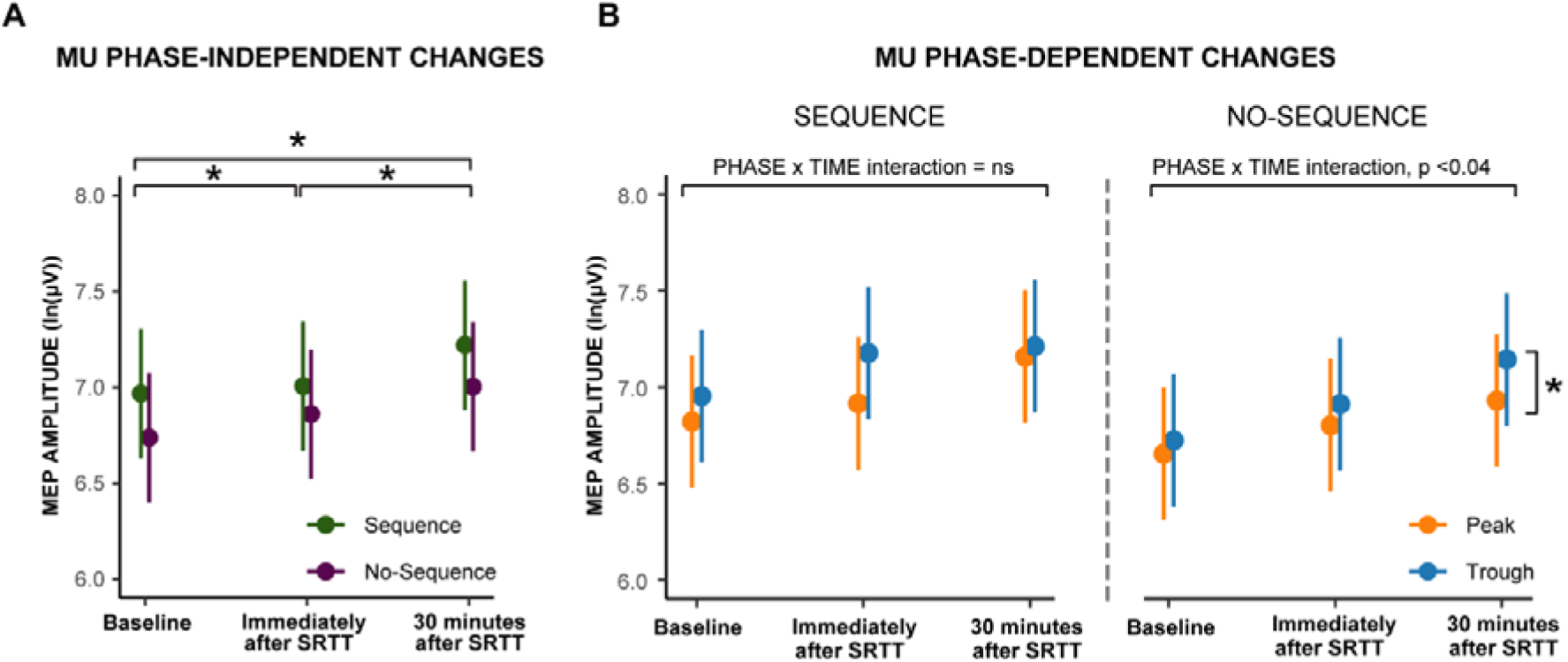
Mu phase-independent and phase-dependent changes in MEP amplitudes. **A)** Group-level linear mixed-effects model coefficients depicting relationships between mu phase-independent MEP amplitudes and group at baseline, immediately and 30 minutes after SRTT acquisition. **B)** Group-level linear mixed-effects model coefficients depicting relationships between mu phase-dependent MEP amplitudes and group at baseline, immediately, and 30 minutes after SRTT acquisition. For A and B, dots represent model coefficients and error bars indicate SEM. For A, black asterisks reflect significant post hoc pairwise comparisons across time points. For B, text above each panel indicates results of the PHASE x TIME interaction, and the black asterisk reflects significant GROUP x PHASE x TIME interaction for the no-sequence group.

Additional post hoc pairwise comparisons showed no differences between peak and trough MEP amplitudes at baseline (β = –0.08, t ratio = –1.68, p= 0.33), but MEP amplitudes were larger during trough than peak phases 30 minutes after SRTT acquisition (β = –0.22, t ratio = –4.48, p < 0.0001).

To visualize differences in mu phase-dependent MEP amplitude changes between groups (i.e., in the sequence group relative to the no-sequence group) identified by the GROUP x PHASE x TIME interaction, we subtracted no-sequence group model-fitted MEP amplitudes from sequence group model-fitted MEP amplitudes (Figure 6). This subtraction illustrated that sequence-specific learning-related MEP amplitude changes were significantly greater during mu rhythm peak than trough phases 30 minutes after SRTT acquisition.

**Figure 6.**
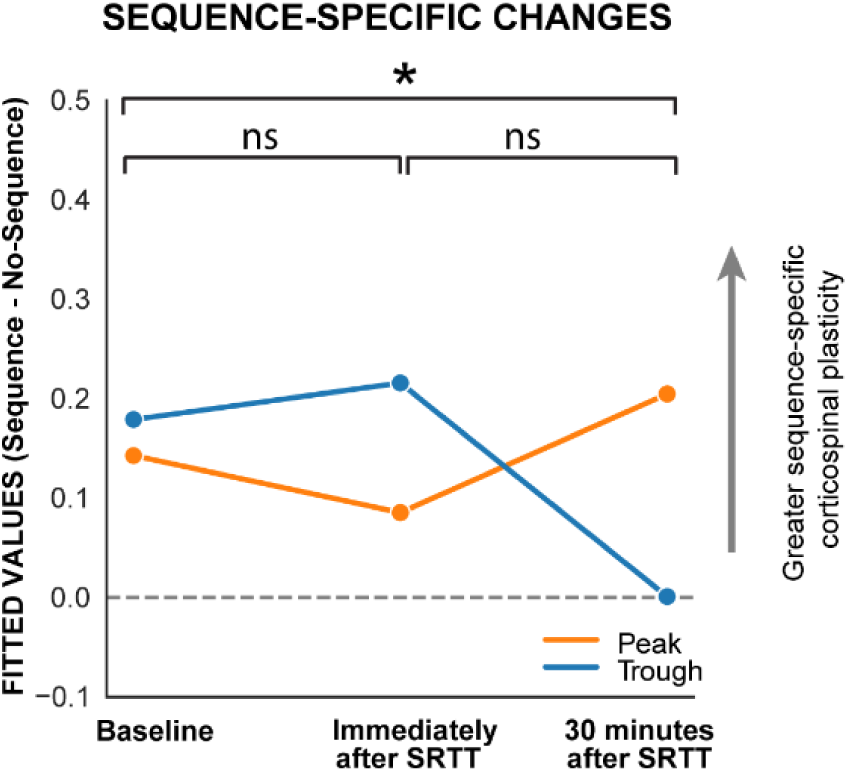
Effects of motor sequence-specific learning on mu phase-dependent MEP amplitude changes. At each time point, group-level model fitted MEP amplitudes obtained during mu peak and trough phases in the no-sequence group were subtracted from corresponding group-level model fitted MEP amplitudes in the sequence group. Black asterisk reflects significant GROUP x PHASE x TIME interaction.

### Relationships between phase-dependent MEP amplitude changes and SRTT performance

Finally, we examined relationships between mu phase-dependent MEP amplitude changes and SRTT performance. When evaluating relationships between peak-specific MEP amplitude changes and SRTT performance, there was a significant effect of GROUP (F = 9.07, p = .005), MEP ratio (F = 8.32, p = .006), and a significant GROUP x MEP ratio interaction (F = 8.82, p = .005), but no main effect of TASK (F = 0.06, p = .8) nor a GROUP x MEP ratio x TASK interaction (F = 0.01, p = .9). The GROUP x MEP ratio interaction was driven by a significantly stronger negative relationship between peak-specific MEP amplitude changes and SRTT performance for the sequence group relative to the no-sequence group (Figure 7A). In contrast, when evaluating relationships between trough-specific MEP amplitude changes and SRTT performance, there was no main effect of GROUP (F = 1.69, p = .2), MEP ratio (F = 1.7, p = .2), or TASK (F = 1.86, p = .18), nor a GROUP x MEP ratio interaction (F = 1.58, p = .21) or a GROUP x MEP ratio x TASK interaction (F = 0.55, p = .46). That is, the relationship between trough-specific MEP amplitude changes and SRTT performance did not significantly differ between groups (Figure 7B). Overall, these findings demonstrate that peak-but not trough-specific MEP amplitude changes were significantly more negatively associated with SRTT performance in the sequence group than in the no-sequence group.

**Figure 7.**
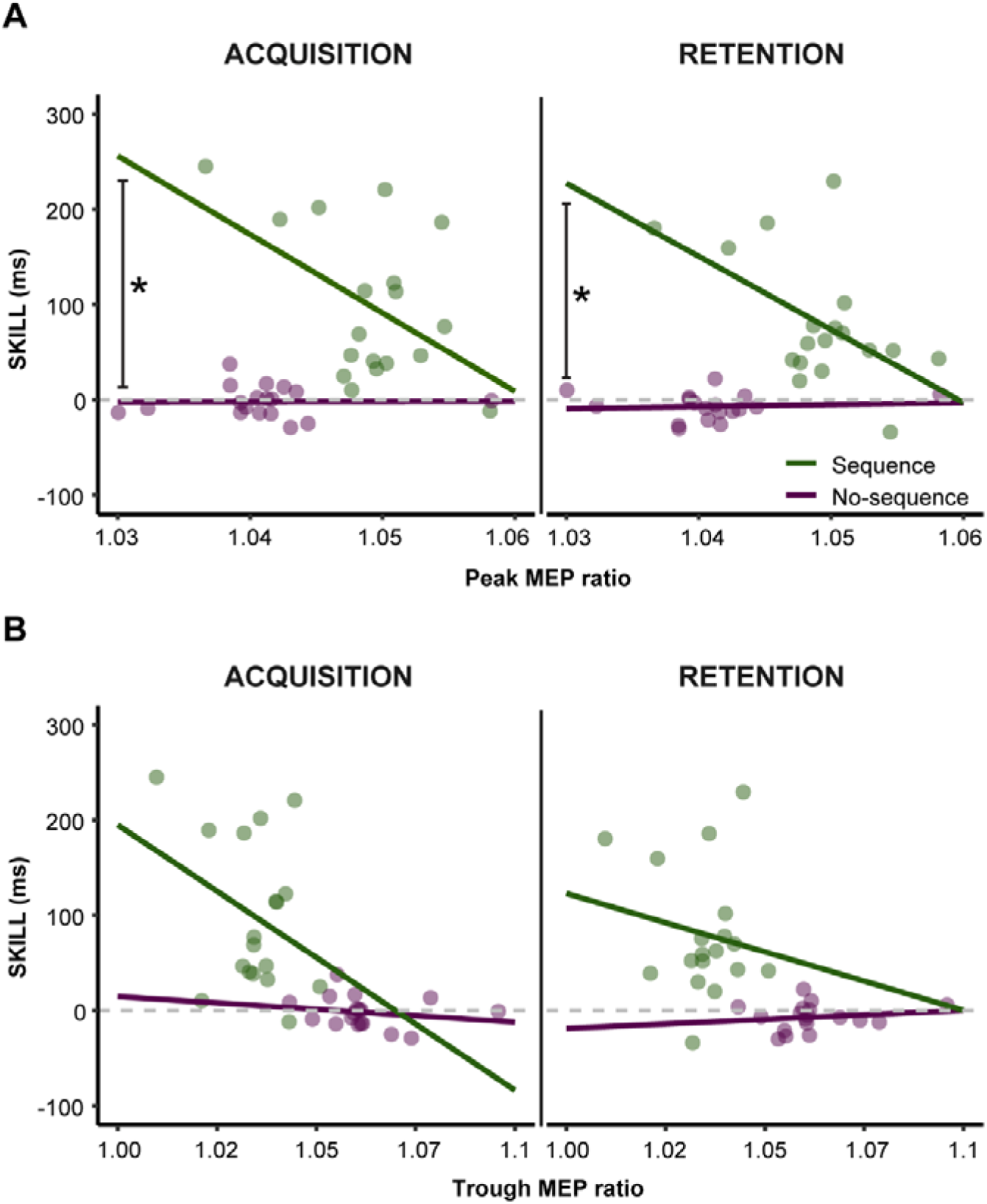
Relationships between mu phase-dependent MEP amplitude changes and SRTT performance. **(A)** Relationship between mu peak-specific MEP amplitude changes and SRTT acquisition skill (left) and retention skill (right). **(B)** Relationships between trough-specific MEP amplitude changes and SRTT acquisition skill (left) and retention skill (right). Phase-specific MEP amplitude changes are represented as the ratio between model-fitted phase-specific MEPs obtained 30 minutes after SRTT acquisition and those obtained at baseline. Larger MEP ratio values indicate greater peak-specific (A) or trough-specific (B) MEP amplitude changes. For both A and B, larger skill values indicate greater reaction time improvements during SRTT acquisition (left) and retention (right). Dots represent individual participant data and lines represent the best fit line. For A, black asterisks reflect significant GROUP x MEP ratio interaction.

## Discussion

Based on our previous work showing that mu phase-dependent TMS interventions coupled to sensorimotor mu rhythm trough but not peak phases improve motor sequence learning, we predicted that motor sequence learning-related corticospinal plasticity would be more evident when measured during sensorimotor mu rhythm trough than peak phases. To test this hypothesis, we measured MEP amplitudes during mu rhythm peak, trough, and random phases immediately before and after healthy adults performed a version of the implicit SRTT that included either an embedded, repeating motor sequence (sequence group) or no sequence (no-sequence group). Behaviorally, participants in the sequence group acquired greater sequence-specific knowledge than participants in the no-sequence group and retained this knowledge over time. Physiologically, participants in both groups showed mu phase-independent increases in corticospinal output following SRTT acquisition, reflecting the occurrence of significant mu phase-independent corticospinal plasticity that did not differ between participants who did and did not acquire sequence-specific knowledge. However, we identified differences in mu phase-dependent changes in corticospinal output between groups. In the sequence group, SRTT acquisition elicited corticospinal plasticity that was similar in magnitude across mu peak and trough phases. In contrast, in the no-sequence group, SRTT acquisition elicited corticospinal plasticity that was greater during mu trough than peak phases. Direct comparisons between groups showed that participants in the sequence group exhibited greater peak-specific corticospinal plasticity than those in the no-sequence group. Further, the magnitude of peak-but not trough-specific corticospinal plasticity was significantly more negatively associated with the magnitude of sequence-specific learning for participants in the sequence relative to the no-sequence group. These findings strongly suggest that performance of each SRTT variant elicits phase-independent corticospinal plasticity, while sequence-specific learning elicits stronger corticospinal plasticity during mu rhythm peak than trough phases.

Motor learning potentiates corticospinal output (Pascual-Leone et al., 1994; Perez et al., 2007) and occludes subsequent induction of LTP-like corticospinal plasticity (Cantarero et al., 2013a; Cantarero et al., 2013b). This occlusion is thought to support retention of newly-acquired motor skills, such that its experimental reversal significantly diminishes retention (Cantarero et al., 2013a). Thus, LTP-like corticospinal plasticity and its occlusion both play key mechanistic roles in motor learning. Like motor learning, repetitive TMS interventions can also induce corticospinal plasticity. Specifically, recent studies using real-time EEG-TMS have shown that both low– and high-frequency repetitive TMS interventions preferentially induce LTP-like corticospinal plasticity when applied during sensorimotor mu trough but not peak phases (Baur et al., 2020; Zrenner et al., 2018). Such mu phase-dependent corticospinal plasticity could therefore reflect a common neurophysiological mechanism underlying the effects of repetitive TMS interventions and motor learning. If mu phase-dependent corticospinal plasticity does support motor learning, then TMS interventions should more strongly affect motor learning when delivered during specific mu phases. We previously evaluated this possibility by measuring the effects of a mu phase-dependent repetitive TMS intervention on offline changes in explicit motor sequence task performance (Hussain et al., 2021). In that study, a repetitive TMS intervention delivered during mu trough but not peak phases enhanced offline gains in motor sequence performance. These findings raised the intriguing possibility that the neurophysiological mechanisms underlying motor learning may be most active and thus most susceptible to modification during sensorimotor mu trough phases. However, in the current study, sequence-specific learning elicited mu phase-dependent corticospinal plasticity that was stronger during mu peak than trough phases.

What could explain these surprising findings? Our experimental design directly compared changes in MEP amplitudes following performance of one of two related motor tasks. In the sequence group, participants performed repeated keypresses while also acquiring knowledge of an embedded sequence, while in the no-sequence group, participants performed repeated keypresses in the absence of any embedded sequence. To isolate the effects of sequence learning on mu phase-dependent corticospinal plasticity, we therefore subtracted no-sequence group model-fitted MEP amplitudes from sequence group model-fitted MEP amplitudes. This direct contrast revealed that sequence learning, after accounting for the effects of repeated keypresses, induced mu phase-dependent corticospinal plasticity that was stronger during mu peak than trough phases. Consistent with this finding, we also observed a significant negative relationship between peak-specific corticospinal plasticity and sequence-specific learning. We previously proposed that mu rhythm peak phases reflect a neutral excitability state that is not strongly affected by synchronization of corticothalamic inputs to layer V pyramidal cells (Hussain et al., 2019). Specifically, M1 single neuron spiking rates are lower during mu rhythm peak than trough phases in non-human primates (Haegens et al., 2011), which may correspond to the decreased corticospinal output observed during mu peak relative to trough phases in humans (Bergmann et al., 2019; Hussain et al., 2019; Suresh & Hussain, 2023). Indeed, Bergmann et al. (2019) showed that mu peak phases reflect a lack of pulsed excitation rather than pulsed inhibition of corticospinal output. Together, these previous findings suggest that mu peaks reflect brief quiescent windows during which corticospinal and sensorimotor network communication is diminished relative to mu troughs. As a result, neural activity should only weakly propagate through an extended sensorimotor network during mu peak phases. We and others previously provided empirical support for this possibility by showing that (1) TMS-related whole-brain broadband power, an indirect marker of population-level neuronal spiking (Gao et al., 2017; Manning et al., 2009), is weaker when M1 TMS delivery coincides with mu rhythm peak than trough phases (Hussain et al., 2021), and (2) interhemispheric communication between bilateral motor cortices is lower during mu peak than trough phases (Stefanou et al., 2018). The decreased network communication that occurs during mu peak phases may prevent motor sequence memories from being disrupted by task-irrelevant sensory information carried by thalamocortical projections, such that sequence memories can be replayed and consolidated by sensorimotor and hippocampal networks without interference (Buch et al., 2021; Jacobacci et al., 2020; Robertson & Genzel, 2020). This idea aligns with previous work showing that wakeful rest after motor sequence learning supports consolidation and retention of the learned sequence (Bönstrup et al., 2019, 2020; Buch et al., 2021; Robertson, 2019; Robertson et al., 2004; Walker et al., 2003). Mu peak phases could therefore provide a protected time window during which replay and consolidation of motor sequence memories may occur. Such a scenario could explain why motor sequence-specific learning-related corticospinal plasticity was more pronounced during mu peak phases in the current study. Further, we also observed a negative relationship between peak-specific MEP amplitude changes and SRTT performance that was significantly stronger in the sequence group than the no-sequence group. Although this result may at first appear counterintuitive, it is possible that poorer motor sequence-specific learning more strongly recruits peak-specific corticospinal plasticity processes in order to protect newly-formed yet fragile sequence memories from interference during consolidation. Thus, peak-specific corticospinal plasticity changes could reflect a compensatory, protective mechanism that is most prominent in participants who initially experience the least sequence-specific learning. Future work could directly test the causal nature of the correlative relationship observed in the current study by evaluating the behavioral consequences of experimentally altering mu peak phase properties immediately after motor sequence learning using either neurostimulation or closed-loop neurofeedback paradigms.

Although we previously suggested that the neurophysiological mechanisms underlying motor sequence learning may be most active during sensorimotor mu trough phases (Hussain et al., 2021), here we report that corticospinal plasticity following motor sequence-specific learning is more strongly expressed during mu peak than trough phases. Two key differences between our previous work and the current study could explain these diverging results. First, the current study used the implicit SRTT, while our previous work used the explicit motor sequence learning task. Although both tasks involve acquisition of sequential knowledge, they rely on dissimilar sensorimotor networks. Specifically, the implicit SRTT likely more heavily recruits corticostriatal circuits (Grafton et al., 1995; Hazeltine et al., 1997), while the explicit motor sequence learning task may more strongly engage prefrontal and premotor areas (Pinsard et al., 2019; Yokoi & Diedrichsen, 2019). Given the fundamental role of oscillatory phase in inter-regional neural communication (Fries, 2005, 2015; Womelsdorf et al., 2007), these two brain networks may optimally interact with sensorimotor cortical neuronal populations during different mu rhythm phases. Second, the current study directly assessed within-session phase-dependent neurophysiological changes induced by motor sequence-specific learning, whereas our previous work examined the influence of a phase-dependent TMS intervention on between-session behavioral changes in motor sequence performance. Thus, corticospinal plasticity supporting motor sequence-specific learning may be most evident during mu peak phases, while TMS interventions may best improve motor sequence learning during mu trough phases. Along these lines, it is possible that the neurophysiological mechanisms which protect newly-formed motor sequence memories from interference during consolidation are maximally active during mu peak phases, such that peak-specific TMS interventions cannot further upregulate them. If so, the same protective mechanisms may be only partially active during mu trough phases, allowing them to be further enhanced by trough-specific TMS interventions. Regardless, our current findings and previous work both support the notion that the neurophysiological mechanisms underlying motor sequence learning are sensorimotor mu phase-dependent.

Several strengths of this study exist. First, our experimental design included two groups: one that experienced sequence learning and one that did not. This design allowed us to dissociate the effects of sequence-specific and non-sequence-specific learning on mu phase-independent and mu phase-dependent corticospinal plasticity. Of note, non-specific effects of motor task engagement are often not considered in motor sequence learning studies (Bönstrup et al., 2019, 2020; Perez et al., 2007; Walker et al., 2003), including our own previous work (Hussain et al., 2021). However, without directly contrasting phase-dependent corticospinal plasticity between the sequence and no-sequence groups, the current study would have failed to identify any role of mu phase-dependent corticospinal plasticity in sequence learning. Second, we quantified both mu phase-independent and phase-dependent corticospinal plasticity. This quantification showed that mu phase-independent corticospinal plasticity can be elicited by repeated keypresses even in the absence of sequence learning itself. Third, participants were not required to meet any pre-defined neurophysiological criteria to participate in this study; our results are likely to be more generalizable across diverse groups of participants than previous studies using similar techniques (Baur et al., 2020; Zrenner et al., 2018). Finally, rather than inferring the existence of a phase-dependent learning mechanism from an interventional study (Hussain et al., 2021), here we specifically quantified phase-dependent neurophysiological changes induced by sequence-specific and non-sequence-specific learning. Thus, the results of the current study can be directly attributed to motor sequence-specific learning rather than the combined effects of motor sequence learning and a phase-dependent TMS intervention.

While our study provides novel insights into phase-dependent mechanisms of motor learning, it is not without limitations. First, previous studies have shown that mu rhythm phase and power interdependently influence corticospinal output at rest (Hussain et al., 2019; Ozdemir et al., 2022; Suresh & Hussain, 2023), but here we evaluated only mu phase-dependent changes in corticospinal output. Although this was achieved by introducing a covariate into our statistical models that controlled for mu phase-power interactions, this approach does not capture more complex learning-related changes in mu phase-power interactions. We also did not measure changes in corticospinal output during other mu phases (Wischnewski et al., 2022; Zrenner et al., 2023), other sensorimotor rhythms (Hussain et al., 2019; Wischnewski et al., 2022) or broadband activity (Davis et al., 2022). Experimental designs using phase-independent measures of corticospinal excitability and post hoc trial sorting approaches may be better suited to explore the contribution of other brain rhythms and phases to learning-related corticospinal plasticity.

In conclusion, we report that motor sequence-specific learning elicits mu peak-specific corticospinal plasticity, and that the magnitude of this peak-specific corticospinal plasticity is significantly negatively associated with sequence-specific learning. These findings provide first direct evidence that motor learning recruits mu phase-dependent neurophysiological mechanisms in humans and establish the mechanistic role of sensorimotor mu rhythm phase in behaviorally-relevant corticospinal plasticity.

## Data availability

Data used for analysis are available from the corresponding author upon reasonable request.

## Acknowledgements

We thank Kristen Pulliam for assistance with data collection.

## Author contributions

TS, FI, MZ, MVF and SJH conceived and designed research; TS, MMc, and MMa performed experiments; TS and SJH analyzed data; TS, MVF, and SJH interpreted results of experiments; TS and SJH prepared figures; TS and SJH drafted manuscript; TS, MVF, and SJH edited and revised manuscript, TS, FI, MZ, MMc, MMa, MVF and SJH approved final version of manuscript.

## References

1. Baumgarten, T. J., Schnitzler, A., & Lange, J. (2015). Beta oscillations define discrete perceptual cycles in the somatosensory domain. Proceedings of the National Academy of Sciences, 112(39), 12187–12192. 10.1073/pnas.1501438112

2. Baur, D., Galevska, D., Hussain, S., Cohen, L. G., Ziemann, U., & Zrenner, C. (2020). Induction of LTD-like corticospinal plasticity by low-frequency rTMS depends on pre-stimulus phase of sensorimotor μ-rhythm. Brain Stimulation, 13(6), 1580–1587. 10.1016/j.brs.2020.09.005

3. Bergmann, T. O., Lieb, A., Zrenner, C., & Ziemann, U. (2019). Pulsed Facilitation of Corticospinal Excitability by the Sensorimotor μ-Alpha Rhythm. Journal of Neuroscience, 39(50), 10034–10043. 10.1523/JNEUROSCI.1730-19.2019

4. Bigoni, C., Pagnamenta, S., Cadic-Melchior, A., Bevilacqua, M., Harquel, S., Raffin, E., & Hummel, F. C. (2024). MEP and TEP features variability: Is it just the brain-state? Journal of Neural Engineering. 10.1088/1741-2552/ad1dc2

5. Bönstrup, M., Iturrate, I., Hebart, M. N., Censor, N., & Cohen, L. G. (2020). Mechanisms of offline motor learning at a microscale of seconds in large-scale crowdsourced data. Npj Science of Learning, 5(1), Article 1. 10.1038/s41539-020-0066-9

6. Bönstrup, M., Iturrate, I., Thompson, R., Cruciani, G., Censor, N., & Cohen, L. G. (2019). A Rapid Form of Offline Consolidation in Skill Learning. Current Biology, 29(8), 1346–1351.e4. 10.1016/j.cub.2019.02.049

7. Borckardt, J. J., Nahas, Z., Koola, J., & George, M. S. (2006). Estimating Resting Motor Thresholds in Transcranial Magnetic Stimulation Research and Practice: A computer Simulation Evaluation of Best Methods. The Journal of ECT, 22(3), 169. 10.1097/01.yct.0000235923.52741.72

8. Buch, E. R., Claudino, L., Quentin, R., Bönstrup, M., & Cohen, L. G. (2021). Consolidation of human skill linked to waking hippocampo-neocortical replay. Cell Reports, 35(10). 10.1016/j.celrep.2021.109193

9. Busch, N. A., Dubois, J., & VanRullen, R. (2009). The Phase of Ongoing EEG Oscillations Predicts Visual Perception. Journal of Neuroscience, 29(24), 7869–7876. 10.1523/JNEUROSCI.0113-09.2009

10. Busch, N. A., & VanRullen, R. (2010). Spontaneous EEG oscillations reveal periodic sampling of visual attention. Proceedings of the National Academy of Sciences, 107(37), 16048–16053. 10.1073/pnas.1004801107

11. Cantarero, G., Lloyd, A., & Celnik, P. (2013). Reversal of Long-Term Potentiation-Like Plasticity Processes after Motor Learning Disrupts Skill Retention. Journal of Neuroscience, 33(31), 12862–12869. 10.1523/JNEUROSCI.1399-13.2013

12. Cantarero, G., Tang, B., O’Malley, R., Salas, R., & Celnik, P. (2013). Motor Learning Interference Is Proportional to Occlusion of LTP-Like Plasticity. Journal of Neuroscience, 33(11), 4634–4641. 10.1523/JNEUROSCI.4706-12.2013

13. Davis, Z. W., Muller, L., & Reynolds, J. H. (2022). Spontaneous Spiking Is Governed by Broadband Fluctuations. The Journal of Neuroscience, 42(26), 5159–5172. 10.1523/JNEUROSCI.1899-21.2022

14. Destrebecqz, A., Peigneux, P., Laureys, S., Degueldre, C., Fiore, G. D., Aerts, J., Luxen, A., Linden, M. V. D., Cleeremans, A., & Maquet, P. (2005). The neural correlates of implicit and explicit sequence learning: Interacting networks revealed by the process dissociation procedure. Learning & Memory, 12(5), 480–490. 10.1101/lm.95605

15. Donoghue, T., Haller, M., Peterson, E. J., Varma, P., Sebastian, P., Gao, R., Noto, T., Lara, A. H., Wallis, J. D., Knight, R. T., Shestyuk, A., & Voytek, B. (2020). Parameterizing neural power spectra into periodic and aperiodic components. Nature Neuroscience, 23(12), Article 12. 10.1038/s41593-020-00744-x

16. Dugué, L., Marque, P., & VanRullen, R. (2011). The Phase of Ongoing Oscillations Mediates the Causal Relation between Brain Excitation and Visual Perception. Journal of Neuroscience, 31(33), 11889–11893. 10.1523/JNEUROSCI.1161-11.2011

17. Fries, P. (2005). A mechanism for cognitive dynamics: Neuronal communication through neuronal coherence. Trends in Cognitive Sciences, 9(10), 474–480. 10.1016/j.tics.2005.08.011

18. Fries, P. (2015). Rhythms for Cognition: Communication through Coherence. Neuron, 88(1), 220–235. 10.1016/j.neuron.2015.09.034

19. Fuentemilla, L., Penny, W. D., Cashdollar, N., Bunzeck, N., & Düzel, E. (2010). Theta-Coupled Periodic Replay in Working Memory. Current Biology, 20(7), 606–612. 10.1016/j.cub.2010.01.057

20. Gao, R., Peterson, E. J., & Voytek, B. (2017). Inferring synaptic excitation/inhibition balance from field potentials. NeuroImage, 158, 70–78. 10.1016/j.neuroimage.2017.06.078

21. Grafton, S. T., Hazeltine, E., & Ivry, R. (1995). Functional Mapping of Sequence Learning in Normal Humans. Journal of Cognitive Neuroscience, 7(4), 497–510. 10.1162/jocn.1995.7.4.497

22. Haegens, S., Nácher, V., Luna, R., Romo, R., & Jensen, O. (2011). α-Oscillations in the monkey sensorimotor network influence discrimination performance by rhythmical inhibition of neuronal spiking. Proceedings of the National Academy of Sciences, 108(48), 19377–19382. 10.1073/pnas.1117190108

23. Hasselmo, M. E., Bodelón, C., & Wyble, B. P. (2002). A Proposed Function for Hippocampal Theta Rhythm: Separate Phases of Encoding and Retrieval Enhance Reversal of Prior Learning. Neural Computation, 14(4), 793–817. 10.1162/089976602317318965

24. Hazeltine, E., Grafton, S. T., & Ivry, R. (1997). Attention and stimulus characteristics determine the locus of motor-sequence encoding. A PET study. Brain, 120(1), 123–140. 10.1093/brain/120.1.123

25. Hjorth, B. (1975). An on-line transformation of EEG scalp potentials into orthogonal source derivations. Electroencephalography and Clinical Neurophysiology, 39(5), 526–530. 10.1016/0013-4694(75)90056-5

26. Hussain, S. J., Claudino, L., Bönstrup, M., Norato, G., Cruciani, G., Thompson, R., Zrenner, C., Ziemann, U., Buch, E., & Cohen, L. G. (2019). Sensorimotor Oscillatory Phase– Power Interaction Gates Resting Human Corticospinal Output. Cerebral Cortex, 29(9), 3766–3777. 10.1093/cercor/bhy255

27. Hussain, S. J., Cohen, L. G., & Bönstrup, M. (2019). Beta rhythm events predict corticospinal motor output. Scientific Reports, 9(1), 18305. 10.1038/s41598-019-54706-w

28. Hussain, S. J., Vollmer, M. K., Stimely, J., Norato, G., Zrenner, C., Ziemann, U., Buch, E. R., & Cohen, L. G. (2021). Phase-dependent offline enhancement of human motor memory. Brain Stimulation, 14(4), 873–883. 10.1016/j.brs.2021.05.009

29. Jacobacci, F., Armony, J. L., Yeffal, A., Lerner, G., Amaro, E., Jovicich, J., Doyon, J., & Della-Maggiore, V. (2020). Rapid hippocampal plasticity supports motor sequence learning. Proceedings of the National Academy of Sciences, 117(38), 23898–23903. 10.1073/pnas.2009576117

30. Kerrén, C., Linde-Domingo, J., Hanslmayr, S., & Wimber, M. (2018). An Optimal Oscillatory Phase for Pattern Reactivation during Memory Retrieval. Current Biology, 28(21), 3383–3392.e6. 10.1016/j.cub.2018.08.065

31. Kothe, C. (2014). Lab streaming layer (LSL).

32. Manning, J. R., Jacobs, J., Fried, I., & Kahana, M. J. (2009). Broadband Shifts in Local Field Potential Power Spectra Are Correlated with Single-Neuron Spiking in Humans. Journal of Neuroscience, 29(43), 13613–13620. 10.1523/JNEUROSCI.2041-09.2009

33. Nissen, M. J., & Bullemer, P. (1987). Attentional requirements of learning: Evidence from performance measures. Cognitive Psychology, 19(1), 1–32. 10.1016/0010-0285(87)90002-8

34. Oldfield, R. C. (1971). The assessment and analysis of handedness: The Edinburgh inventory. Neuropsychologia, 9(1), 97–113. 10.1016/0028-3932(71)90067-4

35. Oostenveld, R., Fries, P., Maris, E., & Schoffelen, J.-M. (2010). FieldTrip: Open Source Software for Advanced Analysis of MEG, EEG, and Invasive Electrophysiological Data. Computational Intelligence and Neuroscience, 2011, e156869. 10.1155/2011/156869

36. Ozdemir, R. A., Kirkman, S., Magnuson, J. R., Fried, P. J., Pascual-Leone, A., & Shafi, M. M. (2022). Phase matters when there is power: Phasic modulation of corticospinal excitability occurs at high amplitude sensorimotor mu-oscillations. Neuroimage: Reports, 2(4), 100132. 10.1016/j.ynirp.2022.100132

37. Pascual-Leone, A., Grafman, J., & Hallett, M. (1994). Modulation of cortical motor output maps during development of implicit and explicit knowledge. Science, 263(5151), 1287–1289. 10.1126/science.8122113

38. Perez, M. A., Wise, S. P., Willingham, D. T., & Cohen, L. G. (2007). Neurophysiological Mechanisms Involved in Transfer of Procedural Knowledge. Journal of Neuroscience, 27(5), 1045–1053. 10.1523/JNEUROSCI.4128-06.2007

39. Pinsard, B., Boutin, A., Gabitov, E., Lungu, O., Benali, H., & Doyon, J. (2019). Consolidation alters motor sequence-specific distributed representations. eLife, 8, e39324. 10.7554/eLife.39324

40. Robertson, E. M. (2019). Skill Memory: Mind the Ever-Decreasing Gap for Offline Processing. Current Biology, 29(8), R287–R289. 10.1016/j.cub.2019.03.007

41. Robertson, E. M., & Genzel, L. (2020). Memories replayed: Reactivating past successes and new dilemmas. Philosophical Transactions of the Royal Society B: Biological Sciences, 375(1799), 20190226. 10.1098/rstb.2019.0226

42. Robertson, E. M., Pascual-Leone, A., & Miall, R. C. (2004). Current concepts in procedural consolidation. Nature Reviews Neuroscience, 5(7), 576–582. 10.1038/nrn1426

43. Shirinpour, S., Alekseichuk, I., Mantell, K., & Opitz, A. (2020). Experimental evaluation of methods for real-time EEG phase-specific transcranial magnetic stimulation. Journal of Neural Engineering, 17(4), 046002. 10.1088/1741-2552/ab9dba

44. Siegel, M., Warden, M. R., & Miller, E. K. (2009). Phase-dependent neuronal coding of objects in short-term memory. Proceedings of the National Academy of Sciences, 106(50), 21341–21346. 10.1073/pnas.0908193106

45. Stefanou, M.-I., Desideri, D., Belardinelli, P., Zrenner, C., & Ziemann, U. (2018). Phase Synchronicity of μ-Rhythm Determines Efficacy of Interhemispheric Communication Between Human Motor Cortices. The Journal of Neuroscience, 38(49), 10525– 10534. 10.1523/JNEUROSCI.1470-18.2018

46. Suresh, T., & Hussain, S. J. (2023a). Re-evaluating the contribution of sensorimotor mu rhythm phase and power to human corticospinal output: A replication study. Brain Stimulation. 10.1016/j.brs.2023.05.022

47. Suresh, T., & Hussain, S. J. (2023b). Re-evaluating the contribution of sensorimotor mu rhythm phase and power to human corticospinal output: A replication study. *Brain Stimulation: Basic*, Translational, and Clinical Research in Neuromodulation, 16(3), 936–938. 10.1016/j.brs.2023.05.022

48. Tunovic, S., Press, D. Z., & Robertson, E. M. (2014). A Physiological Signal That Prevents Motor Skill Improvements during Consolidation. Journal of Neuroscience, 34(15), 5302–5310. 10.1523/JNEUROSCI.3497-13.2014

49. VanRullen, R., Busch, N. A., Drewes, J., & Dubois, J. (2011). Ongoing EEG Phase as a Trial-by-Trial Predictor of Perceptual and Attentional Variability. Frontiers in Psychology, 2, 60. 10.3389/fpsyg.2011.00060

50. Walker, M. P., Brakefield, T., Allan Hobson, J., & Stickgold, R. (2003). Dissociable stages of human memory consolidation and reconsolidation. Nature, 425(6958), 616–620. 10.1038/nature01930

51. Watrous, A. J., Fell, J., Ekstrom, A. D., & Axmacher, N. (2015). More than spikes: Common oscillatory mechanisms for content specific neural representations during perception and memory. Current Opinion in Neurobiology, 31, 33–39. 10.1016/j.conb.2014.07.024

52. Wilkinson, L., & Jahanshahi, M. (2007). The striatum and probabilistic implicit sequence learning. Brain Research, 1137, 117–130. 10.1016/j.brainres.2006.12.051

53. Wilkinson, L., Khan, Z., & Jahanshahi, M. (2009). The role of the basal ganglia and its cortical connections in sequence learning: Evidence from implicit and explicit sequence learning in Parkinson’s disease. Neuropsychologia, 47(12), 2564–2573. 10.1016/j.neuropsychologia.2009.05.003

54. Wilkinson, L., & Shanks, D. R. (2004). Intentional Control and Implicit Sequence Learning. *Journal of Experimental Psychology: Learning*, Memory, and Cognition, 30(2), 354–369. 10.1037/0278-7393.30.2.354

55. Wilkinson, L., Steel, A., Mooshagian, E., Zimmermann, T., Keisler, A., Lewis, J. D., & Wassermann, E. M. (2015). Online feedback enhances early consolidation of motor sequence learning and reverses recall deficit from transcranial stimulation of motor cortex. Cortex, 71, 134–147. 10.1016/j.cortex.2015.06.012

56. Wischnewski, M., Haigh, Z. J., Shirinpour, S., Alekseichuk, I., & Opitz, A. (2022). The phase of sensorimotor mu and beta oscillations has the opposite effect on corticospinal excitability. *Brain Stimulation: Basic*, Translational, and Clinical Research in Neuromodulation, 15(5), 1093–1100. 10.1016/j.brs.2022.08.005

57. Womelsdorf, T., Schoffelen, J.-M., Oostenveld, R., Singer, W., Desimone, R., Engel, A. K., & Fries, P. (2007). Modulation of Neuronal Interactions Through Neuronal Synchronization. Science, 316(5831), 1609–1612. 10.1126/science.1139597

58. Yokoi, A., & Diedrichsen, J. (2019). Neural Organization of Hierarchical Motor Sequence Representations in the Human Neocortex. Neuron, 103(6), 1178–1190.e7. 10.1016/j.neuron.2019.06.017

59. Zrenner, C., Desideri, D., Belardinelli, P., & Ziemann, U. (2018). Real-time EEG-defined excitability states determine efficacy of TMS-induced plasticity in human motor cortex. Brain Stimulation, 11(2), 374–389. 10.1016/j.brs.2017.11.016

60. Zrenner, C., Kozák, G., Schaworonkow, N., Metsomaa, J., Baur, D., Vetter, D., Blumberger, D. M., Ziemann, U., & Belardinelli, P. (2023). Corticospinal excitability is highest at the early rising phase of sensorimotor µ-rhythm. NeuroImage, 266, 119805. 10.1016/j.neuroimage.2022.119805

